# TopEnzyme: A framework and database for structural coverage of the functional enzyme space

**DOI:** 10.1101/2022.06.13.495871

**Authors:** Karel van der Weg, Holger Gohlke

**Affiliations:** John von Neumann Institute for Computing (NIC), Jülich Supercomputing Centre (JSC), and Institute of Bio- and Geosciences (IBG-4: Bioinformatics), Forschungszentrum Jülich GmbH, 52425 Jülich, Germany; Institute for Pharmaceutical and Medicinal Chemistry, Heinrich Heine University Düsseldorf, 40225 Düsseldorf, Germany

**Author notes:** Corresponding Author: Holger Gohlke, Address: Wilhelm-Johnen-Str., 52425 Jülich, Germany.

**Keywords:** enzyme database, AlphaFold2 comparison, enzyme classification coverage, enzyme structures, enzyme function, function prediction

## Abstract

TopEnzyme is a database of structural enzyme models created with TopModel and is linked to the SWISS-MODEL and AlphaFold Protein Structure Database to provide an overview of structural coverage of the functional enzyme space for over 200,000 enzyme models. It allows the user to quickly obtain representative structural models for 60% of all known enzyme functions. We assessed the models with TopScore and contributed 9039 good-quality and 1297 high-quality structures. Furthermore, we compared these models to AlphaFold2 models with TopScore and found that the TopScore differs only by 0.04 on average in favor of AlphaFold2. We tested TopModel and AlphaFold2 for targets not seen in the respective training databases and found that both methods create qualitatively similar structures. When no experimental structures are available, this database will facilitate quick access to structural models across the currently most extensive structural coverage of the functional enzyme space.

## Background

Recent developments in high-throughput sequencing methods led to a massive increase in sequence information. Databases such as the UniProtKB(1) contain over 225,000,000 sequence records, of which over 550,000 are manually annotated and reviewed in Swiss-Prot. By contrast, the Protein Data Bank (PDB)(2), the worldwide repository of information about the 3D structure of biomolecules, contained 185,539 crystal structures at the end of 2021, of which many are redundant structures, and not all are enzymes. Generally, the topology (fold) of an enzyme is thought to be the major determinant for the given function(3,4). Currently, enzyme function prediction methods often use structures from the PDB for the training data set. However, this can lead to biases in prediction from the protein topology, especially for proteins with a similar topology but a different function, such as TIM-barrels(5) and Rossman folds(6). With recent improvements in protein structure prediction methods(7–10), the availability of high-quality structural models has increased(11). Such structural models will contribute to better coverage and balance of the structural enzyme space.

The currently used databases that categorize structural relationships are SCOP2(12) (Structural Classification of Proteins) and CATH(13) (CATH Protein Structure Classification Database). Both databases provide a detailed and comprehensive description of the structural evolutionary relationships between proteins whose 3D structure has been deposited in the PDB. Although these databases provide information on structural characteristics of the protein and the related functions, further analyses using database-specific classifiers are required to obtain the structural information related to function. While CATH provides FuncFams (functional families), these are not based on the enzymatic functions as classified by the Nomenclature Committee of the International Union of Biochemistry and Molecular Biology (IUBMB)(14). The IUBMB currently curates the list of enzyme commission (EC) numbers. To our knowledge, only the Enzyme Structure Database from the EMBL-EBI relates structures to the enzyme classification. However, this database has not been updated since 2018 and does not cover the newest enzyme class, translocases. In this study, we introduce the database TopEnzyme, where 15,500 representative structural models are categorized by enzyme classification numbers for the currently largest structural coverage of functional enzyme space. TopEnzyme, which includes additional information from SWISS-MODEL(15) and the AlphaFold Protein Structure Database(11) (AlphaFold DB) provides a comprehensive overview and facilitates access to obtain structures associated with specific enzyme functions.

When we started the generation of TopEnzyme, only 22% of the structural enzyme space was covered with respect to the available sequence information in the PDB. Using our deep learning- and template-based software TopModel(7), we generated structural models of 10,125 enzyme domains covering 4,758 different folds, increasing the coverage to 35%. With the release of AlphaFold2(8) and its goal to model the full UniprotKB/Swiss-Prot(11), the current structural coverage of the functional enzyme space is at 60% across SWISS-Model, TopEnzyme, and AlphaFold DB, covering all available sequences with EC annotation in the UniProtKB/Swiss-Prot. We made use of the availability of two complementary protein structure prediction methods to mutually validate structural models and provide a comparison of the structural quality for 2,419 models.

## Construction and Content

Using ExpasyEnzyme(16) (accessed on 12-05-2022), we obtained a complete list of UniprotAC identifiers for 241,125 sequences with an enzyme function annotation according to EC numbers from Swiss-Prot. The first three levels of EC numbers represent the main-, sub-, and subsub-class functions, while the fourth level is the specific enzyme function designation. E.g., the small monomeric GTPase with designation 3.6.5.2 is a hydrolase (3) that acts on acid anhydrides (6) and specifically on guanosine triphosphate (GTP) to facilitate cellular and subcellular movements (5). In total, we find 252 subsub-classes spanning 4926 unique designations. Using MMseqs2(17), we clustered the obtained sequences with an identity cut-off of 30%, such that each cluster represents a homologous cluster in the enzyme fold space(18–20). When a cluster contains more than one subsub-class, we split this cluster into smaller clusters such that we identify a representative for each subsub-class function. For each cluster, we aimed at modeling the representative with TopModel, using templates with a sequence identity > 30% and a sequence coverage > 80%. The refinement procedure in TopModel was skipped as the strict restraints set by the chosen templates should provide structures of sufficient quality while keeping computational costs to a feasible level. This is confirmed for 70 structural models, 10 randomly selected from each enzyme mainclass, which were evaluated as to the effect the refinement procedure has on the TopScore(10) of such models compared to ones without refinement; TopScore is a confidence measure of predicted structural quality. The refinement improved the models by a TopScore of only 0.06 on average (Figure S1). Although TopModel allows for manual template selection, e.g., offering the ability to use templates with bound ligands, this may limit the availability of templates with sufficient quality. Thus, we opted to use all available templates to create more enzyme structural models.

In the case of multi-domain enzymes for which template information is missing for one or more non-catalytic domains, we remove unmodeled regions according to the following criteria: 1) Must be at least ten residues long. 2) Must not contain residues of secondary structure elements (α-helix or β-strand) longer than five residues. 3) Must have a median relative solvent accessible surface area larger than 0.40. 4) Must have a median contact density smaller than four contacts. This prevents the removal of loops close to the binding site. 5) We only remove unmodeled regions from the C- or N-terminal to keep loops within the modeled domain(s). In the case of multi-domain enzymes for which the template contains information on multiple domains, we often model the complete structure.

We created an interactive treemap of the resulting EC space and associated structural models of enzymes (Figure 1, https://cpclab.uni-duesseldorf.de/topenzyme). Each section is mapped by the main-, sub, subsub-class, and designation EC labels. The size of each section represents the proportion of the EC space as given by the number of representative sequences with this EC number. The colors represent the average pLDDT score (8), a confidence measure of predicted structural quality from AlphaFold2, for the representatives of each EC function. The last level represents the designation EC label. There, information is provided on enzymatic function and availability of experimental and representative structure models, as are links to other enzymatic function databases and a download link that generates a bash script to download the representative models from TopEnzyme, SWISS-Model, and AlphaFold DB. This bash script works on all major platforms, provided you have a bash version installed. The meta-data required to create this treemap is available as a csv file in the Supporting Information.

**Figure 1.**
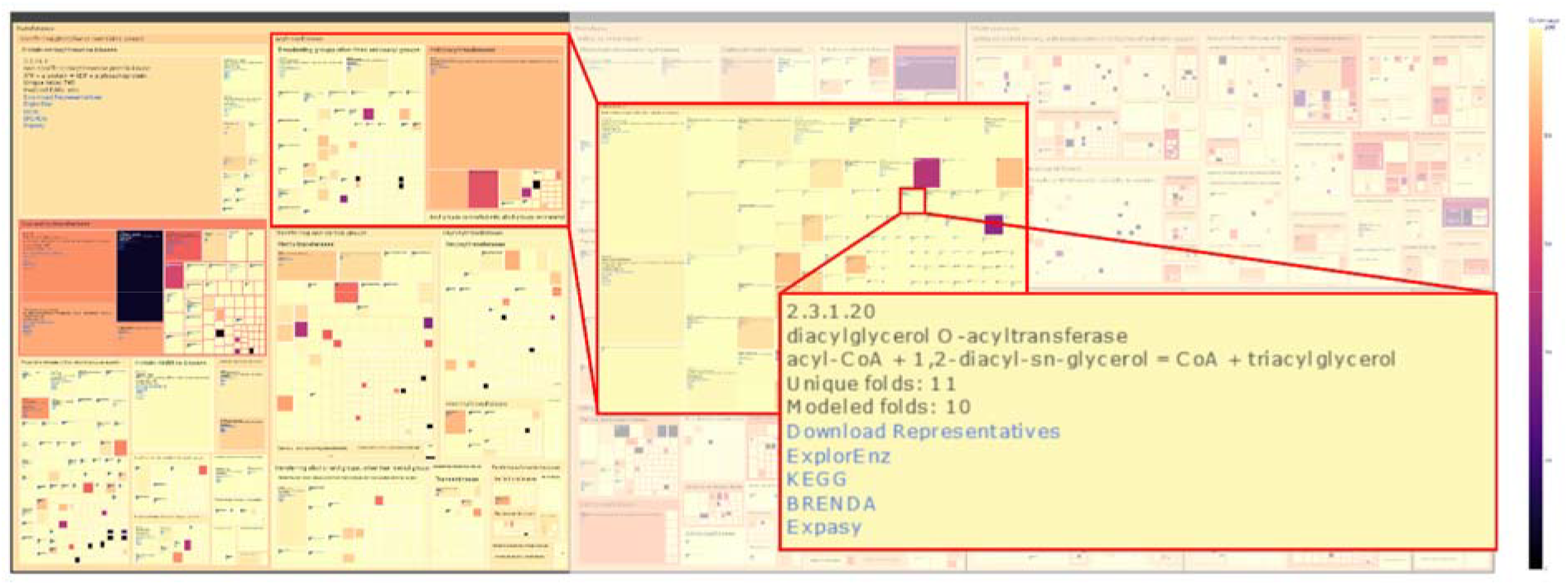
Enzyme map presenting the coverage of EC space with structural models. A screenshot of the interactively explorable enzyme map provided in the Supporting Information (Data S1) and available at https://cpclab.uni-duesseldorf.de/topenzyme/. The color represents the pLDDT for representative models obtained from AlphaFold2 (higher pLDDT numbers indicate better structural quality). The rectangle size represents the number of representative structures for the specific function. The treemap is ordered according to EC classification. By clicking on an area, the next subclass enlarges and shows the information for enzymatic function, enzyme classification, number of (available) folds within this class as determined from sequences in the UniprotKB/Swiss-Prot, a link to the enzyme databases ExplorEnz, KEGG, BRENDA and Expasy, and a download link that generates a script to download all structures available in TopEnzyme, SWISS-MODEL, and AlphaFold DB. This figure is created with plotly.

Using TopScore, we analyzed the quality of the predicted models. 90% of these models are of good quality (TopScore < 0.4), equivalent to 9039 structures spanning 233 subsub-classes (Figure 2a, e). As to secondary structures, TopModel works better for predicting α-helices and β-sheets than loop conformations (Figure 2b). This effect might be caused by bypassing the refinement stage (Figure S1). As expected, the score of our models increases with the sequence similarity of the template, except for models with a sequence similarity > 90% (Figure 2c). Overall, the structural quality of the binding site is similar to that of the entire enzyme. However, we see a larger spread in per-residue scores around the binding site (Figure 2d): Often, secondary structure features in the binding site are of high quality, whereas loop regions contribute to the lower-scoring residues. The exact structure of these loop regions may be less relevant to characterizing the dynamic nature of the protein binding site as these loops have often been shown to move to accommodate the space required for binding the ligand. The spread in binding site model quality might also be due to differences in the structural completeness of the binding sites, e.g., in the case of multi-domain enzymes or allosteric enzymes where activation depends on domain- and ligand binding interactions. Using PocketAnalyzer^PCA^ (21), we determined the average degree of buriedness (DOB) of our binding sites. The majority (77%) of the binding sites are characterized as inside a domain and, hence, are considered complete (Figure 2f). For the binding sites on the surface, we categorize two types: surface and surface (non-complete). The latter fraction (15% of all binding sites) is determined by mapping the binding site location to the homologous template and identifying the presence of not modeled complementary surfaces from global stoichiometry information in the template. By contrast, for the surface fraction (8%), we could not find structural information for complementary interfaces. This does not mean that there is no complementary surface present, just that there is no available information. Each structural model contains the residue-wise TopScore in the B-factor column of the PDB file. This allows the user to investigate the model confidence for specific regions (Figure 2g).

**Fig. 2.**
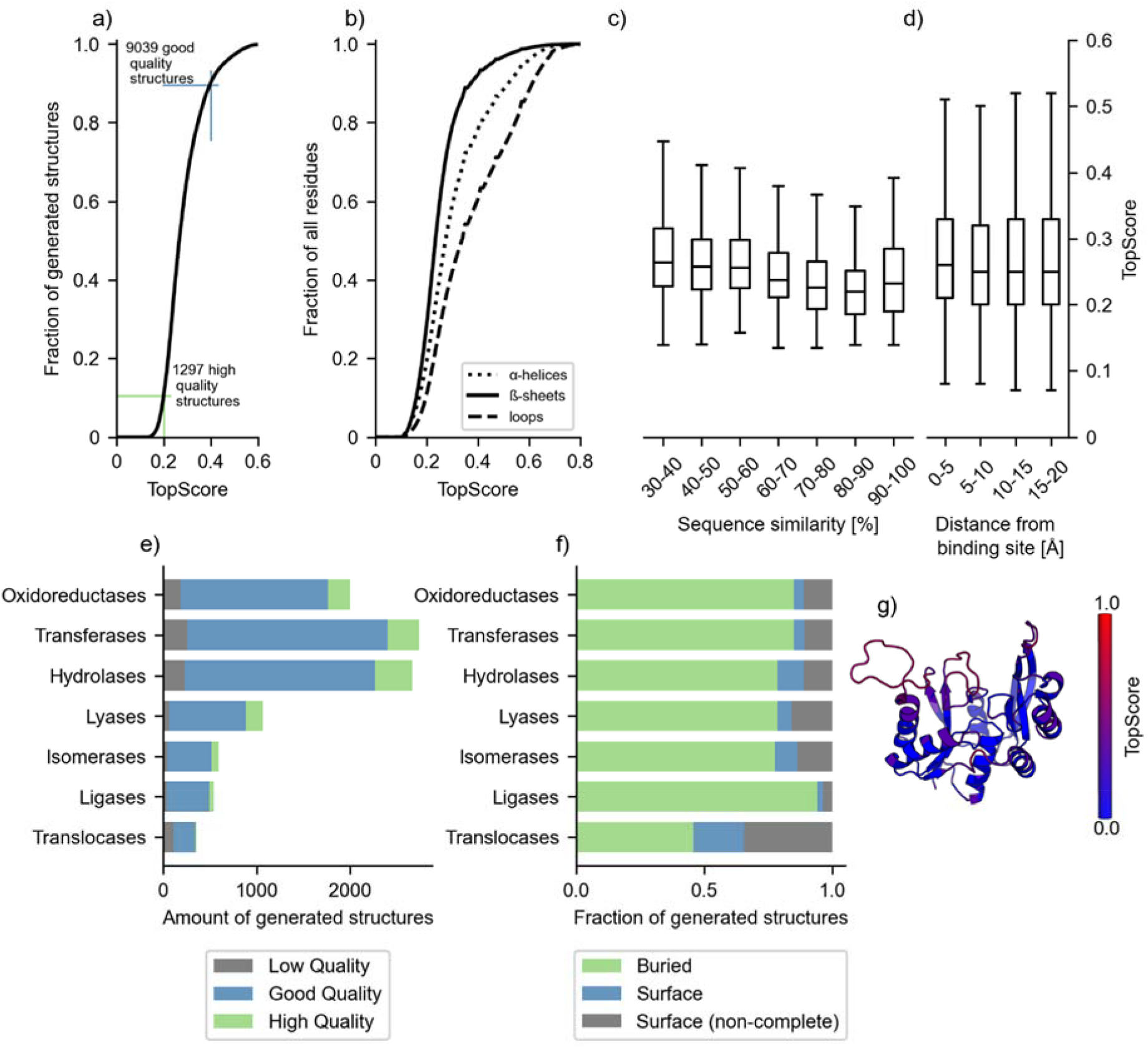
Quality assessment of enzyme models generated with TopModel. **a**, TopScore for all (*n* = 9947) models predicted with TopModel. All models are generated with a template identity > 30% and a coverage > 80%. The green and blue lines indicate the cut-offs for TopScore values associated with high quality (TopScore < 0.2; *n* = 1297) and good quality (TopScore < 0.4; *n* = 9039) structures. TopScore values are bounded between 0 and 1; a lower TopScore is better. **b**, Residue-wise TopScore for all model residues (*n* = 3,191,133) grouped by structural features. The continuous line represents the b-sheet residues (*n* = 629,778), the dash-dot line represents the α-helix residues (*n* = 1,961,309), and the dotted line represents the loop residues (*n* = 600,039). **c**, The distribution of model quality based on the sequence similarity to the template used. The whiskers show the full range of TopScore values. The median is shown on the horizontal notch. **d**, TopScore values for all (*n* = 3,191,133) model residues. The scores are clustered by the distance from the binding site. The boxplot properties are the same as in c. **e**, Coverage of the generated structures by the main enzyme class. The horizontal bars are separated by low (TopScore ≥ 0.4), good (0.2 ≤ TopScore < 0.4), and high (TopScore < 0.2) quality structures. **f**, Fraction of generated structures categorized by the main enzyme class with binding interfaces within a domain (“buried”), binding interfaces on the surface of a domain with (“surface”) and without (“surface (non-complete)”) known complementary domain(s). **g**, An example structure (UniprotAC: Q9CQ28) highlighting the structural per-residue quality as judged by TopScore (see color scale). The image is generated using PyMOL 2.3.0 (PyMOL).

## Utility and Discussion

The intended use of the database is to facilitate research connecting enzyme structure and biochemical function. The database serves this aim with its framework of covering EC space with structural models and easy applicability for users with different levels of expertise (see below). By using familiar identifiers, UniprotACs and EC numbers, and linking to other databases, such as the AlphaFold DB and SWISS-MODEL as well as ExploreEnz, KEGG, BRENDA and Expasy, we provide a framework for comprehensive structural enzyme information linked to enzymatic function.

We envision using generated models over crystal structures important for prediction methods for several reasons. First, some structural noise could make the machine learning method more robust to uncertain information (22,23). Second, proteins are not rigid objects; having a uniform way of generating structural models should be advantageous compared to using experimental structures from different sources or binding states. Third, databases of predicted structural models cover a larger functional enzyme space. Last, this allows for extendibility to information from, e.g., metagenomic approaches, where no structural information is available, but sequences are deposited in the UniProtKB.

Compared to current databases such as the SCOP2 and CATH, our focus is on enzymatic function linked to available enzyme structural models. TopEnzyme starts from enzyme function categorization and provides available structural models from the respective fold space with easy access from the largest collection of generated enzyme models. There are two methods to interact with the database: I) For scientists interested in large-scale analysis, we provide a csv file containing all the meta-data for each UniprotAC identifier. This allows users to download the latest release and incorporate the information into their workflows. II) For scientists interested in a few cases with specific enzyme functions, we provide a visualization in the form of the treemap (Figure 1) hosted on https://cpclab.uni-duesseldorf.de/topenzyme/. The treemap allows users to browse enzymes with specific functions and provides a simple download method to obtain the representative models from the linked databases. In our own project, we used TopEnzyme to quickly obtain representative enzyme structures for building large datasets for a deep learning-based EC number classification.

We plan to update TopEnzyme when there is a new major release to databases of enzyme structural models or structural information for previously unlinked EC classes. As to new features, we intend to integrate a more exhaustive structural data collection from linked databases, as, currently, we collect only the best ranked structural model from each method, while some methods produce ensembles of models. Furthermore, we will improve the search options to include a list of enzymatic functions to move the treemap to the selected function and include structural visualization when selecting a treemap node.

### Comparison to AlphaFold2

To obtain further insights into the quality of our structural models, we compare a proportion of our enzyme structural models with AlphaFold2(8) structures from the AlphaFold DB for the same enzymes. We remove disordered domains on the AlphaFold2 structures in the same way as in ours for fairness. The current AlphaFold2 implementation only folds single domains, although other implementations based on AlphaFold2 have been described that can predict multimers(24,25). We randomly selected 25% in each main class of our database for comparison to AlphaFold2 (Figure 3). While the majority of TopEnzyme structures are available in the AlphaFold DB, TopEnzyme contains few structures unmodeled in the AlphaFold DB yet. We compared TopScore to pLDDT for both regimes (Figure S2). As TopScore and pLDDT predict (1 – lDDT)(8,10) and lDDT(26), both correlate significantly and fairly (*p* < 0.001, *R*^2^ = 0.59). Remarkably, when tested on experimental enzyme structures unseen by both methods (Figure S3), TopScore underestimates and AlphaFold2 overestimates the lDDT. We investigated the majority in the good-quality regime (TopScore values < 0.4; *n* = 1935) and a smaller number in the poor-quality regime (TopScore values ≥ 0.4; *n* = 484) to obtain a comprehensive overview. In the good-qualitative regime, AlphaFold2 performs slightly better than TopModel as judged by TopScore values computed for each model pair, which is consistent across all enzyme main classes. However, in the poor-quality regime, TopModel creates better models than AlphaFold2 for most enzyme classes except transferases and hydrolases. This result suggests that for some target sequences a model created from one or more homologous templates might be better.

**Fig. 3.**
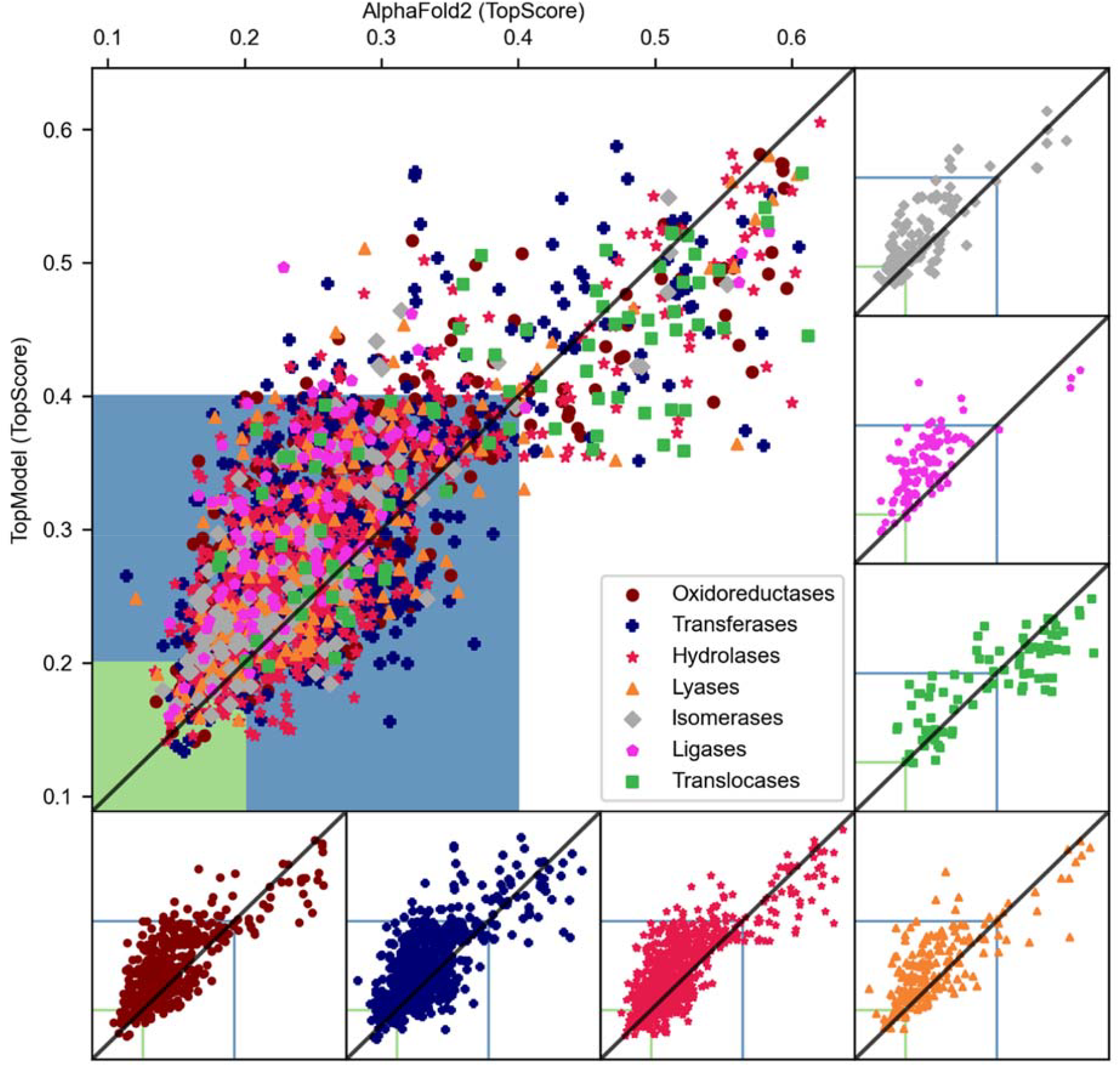
Comparison of structural models generated by TopModel and AlphaFold2. The models generated by TopModel or AlphaFold2 were scored with TopScore. For TopModel, all models are generated with a sequence similarity of > 30% and a coverage of > 80% with respect to the target. The AlphaFold2 models are obtained from the AlphaFold DB, alphafold.ebi.ac.uk. We compared 20% of the models generated by TopModel in the good (TopScore < 0.4; *n* = 1935) and 5% of the models in the bad (TopScore ≥ 0.4; *n* = 484) regime for each enzyme main class against AlphaFold2 structures. The diagonal line represents an equal score between the models. Data points above the diagonal favor AlphaFold2 structures, and data points below the diagonal favor TopModel structures. The blue area represents the score for good-quality models, and the green area represents the score for high-quality models. The panels around the figure show the same content but separated by enzyme main class.

### Comparison to experimental structures

Besides comparing both structure prediction methods, we also compare both methods to recently released X-ray crystallography structures in the PDB (Figure 4). These structures are chosen such that they were not part of the training data for AlphaFold2 or TopModel. In general, both methods produce models of comparable quality, with AlphaFold2 models having a better average TopScore of 0.04.

**Fig. 4.**
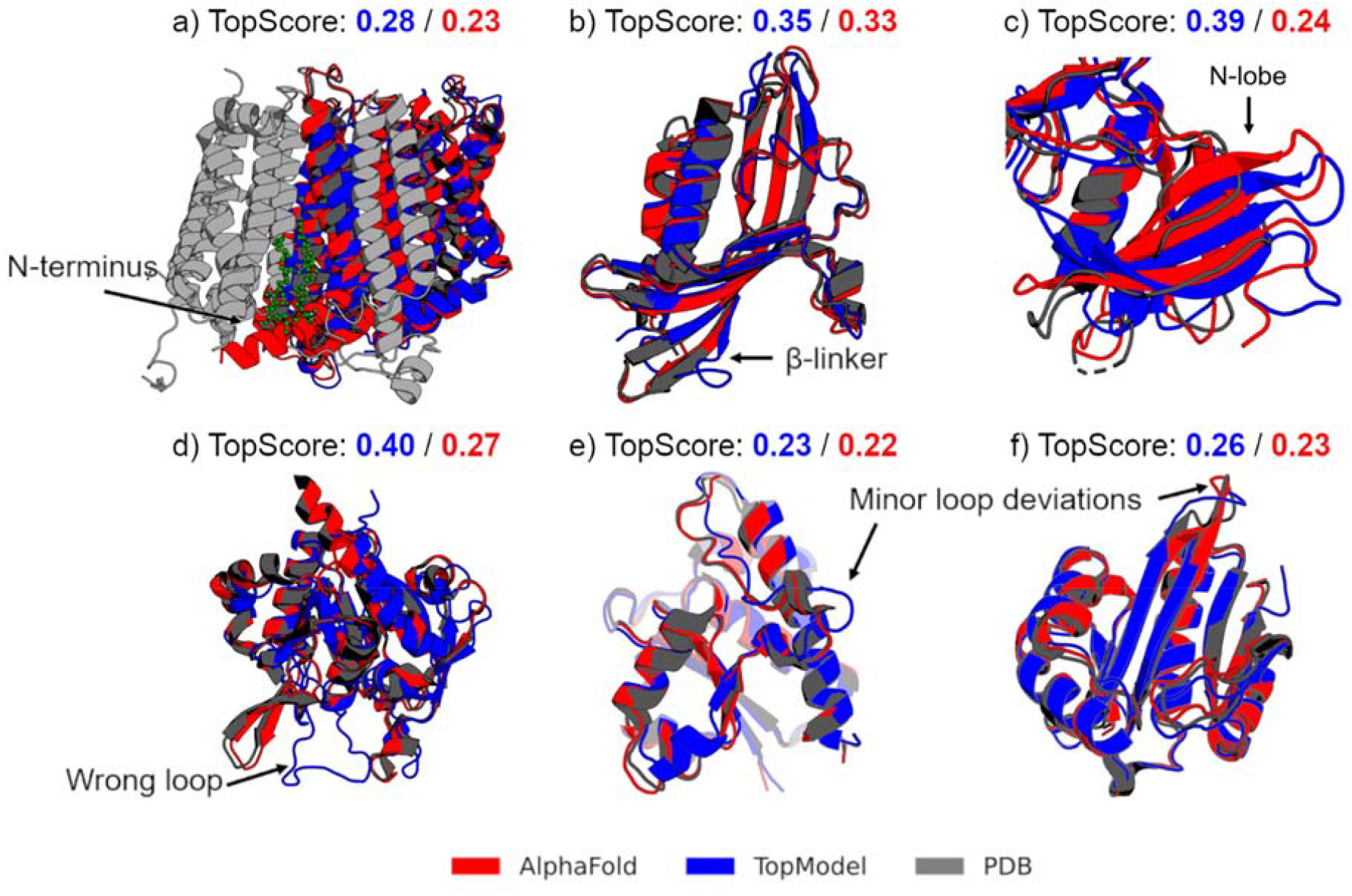
Comparison to experimental structures. An overview of six crystal structures (gray) obtained from the Protein Data Bank compared to AlphaFold2 (red) and TopModel (blue) models. Below the structures, the TopScore for the TopModel and AlphaFold2 models is shown. The arrows correspond to structural features discussed in the text. All structures have been deposited recently and are not present in the training databases for either method. **a**, NADH-ubiquinone oxidoreductase (PDB ID 7A23) in the plant mitochondrial respiratory complex I. In the AlphaFold2 model, an α-helix incorrectly protrudes in the direction of the complementary subunit. **b**, Salicylate 5-hydroxylase (PDB ID 7C8Z). Both methods produce a good-quality structure. **c**, Yeast TFIIK (Kin28/Ccl1/Tfb3) complex (PDB ID 7KUE). Both methods predict a larger b-sheet region that is characterized as mostly coil in the PDB file. **d**, Gamma-glutamyl-gamma-aminobutyrate hydrolase (PDB ID 6VTV). The TopModel model mispredicts part of the fold and creates a random coil region instead of a β-sheet. **e**, Phosphotyrosine protein phosphatase 1 (PDB ID 7CUY). In the TopModel model, a small random coil region diverges from the PDB structure and AlphaFold2 model. **f**, *N*-α-acetyltransferase 30 (PDB ID 7L1K). Both methods create a good-quality structure.

In NADH-ubiquinone oxidoreductase (Figure 4a), the membrane domain is modeled well by both methods, except for the N-terminus, which is uncharacterized in the PDB ID 7A23. In the case of TopModel, this part is modeled as a disordered region, which gets removed by postprocessing. AlphaFold2 predicts this region as an α-helical structure, albeit with low confidence. In both cases, the N-termini stick straight through the binding site for cardiolipin in the crystal structure. This site is recognized as an important site for the stability of the protein domain(27). In Salicylate 5-hydroxylase (Figure 4b), two models were predicted with structural features of excellent quality. However, the loops between the β-strands and the random loops deviate from the crystal structure, which lowers the global score. For the Yeast TFIIK (Kin28/Ccl1/Tfb3) complex (Figure 4c), we focus on the specific N-lobe region of the enzyme(28). The difference in TopScore values between both methods is due to the improved structural features in that domain of the AlphaFold model. However, both methods create a similar deviation in the N-lobe region in that they modeled a larger β-sheet. Note that the crystal structure (PDB ID 7KUE) was refined in conjunction with enzyme CDK2 (PDB ID 1FIN), which is present in both the training database for TopModel and AlphaFold2. If we compare the β-sheet region among the models and CDK2, the models agree perfectly with CDK2(28), where this β-sheet region is much larger. Likely, both methods learned to model this section according to CDK2 instead of predicting the smaller β-sheet seen in Kin28. The TopModel model of Gamma-glutamyl-gamma-aminobutyrate hydrolase (Figure 4d) is an example of insufficient quality. Even though most of the structural features are in good agreement, a loop region should have been modeled as a β-sheet. Finally, for both Phosphotyrosine protein phosphatase (Figure 4e) and *N*-α-acetyltransferase 30 (Figure 4f), the models generated by either method are very good. The TopScore is high, and the structural features are in excellent agreement with the PDB structures (PDB ID 7CUY, PDB ID 7L1K).

Furthermore, we take a detailed look at three binding interfaces for recently released X-ray crystallography structures in the PDB (Figure 5). Only the structural features and loops close to the binding sites are visualized to improve clarity. In the matrix arm for plant mitochondrial respiratory complex I (PDB ID 7A23, Figure 5a), FeS and SF4 clusters are important for the proton pump mechanism in ubiquinone reductase(27,29). For both models, the structural details around the FeS and SF4 clusters are nearly identical to the crystal structure. In the *Mycobacterium tuberculosis* protein FadB2 (PDB ID 6HRD, Figure 5b), the two Rossman folds, β1-α1-β2 and β4-α4-β5, and the α7-α11 region are very well modeled(30). The α2 helix in the TopModel model slightly points away from Coenzyme A. Further investigation of the N-lobe in Yeast TFIIK reveals that for the TopModel model the flexible linker sticks into the ADP binding site and the activation loop deviates from crystal structure (Figure 5c). Yet, predicting these residues exactly as in the crystal structure might be less crucial as they are shown to be the least stable residues of the enzyme(31).

**Fig. 5.**
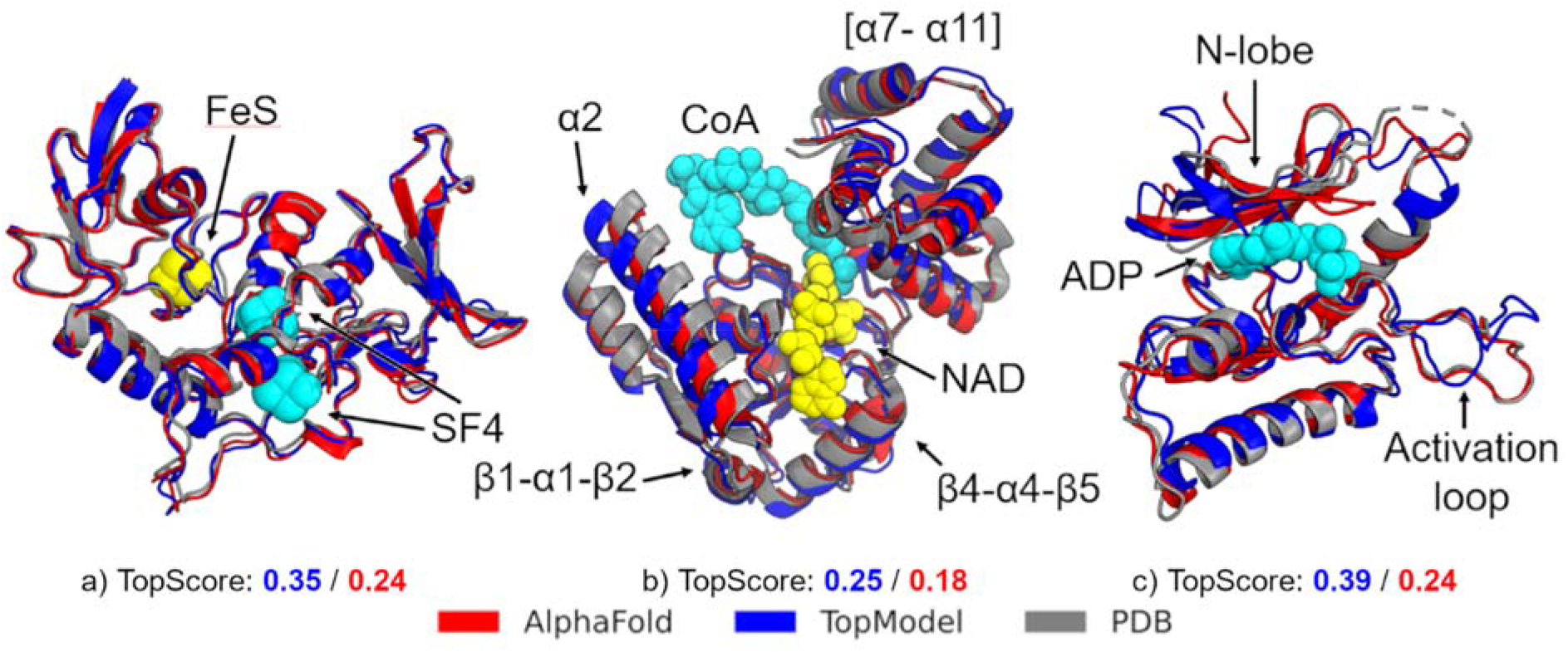
Comparison of binding sites. A cut-out of the binding site for three crystal structures (gray) obtained from the Protein Data Bank and compared to AlphaFold2 (red) and TopModel (blue) models with the corresponding binding ligands. The arrows correspond to structural features discussed in the text. All structures have been deposited recently and are not present in the training databases for either method. Only the structural features close to the binding sites are visualized to improve clarity. **a**, The matrix arm of plant mitochondrial respiratory complex I (PDB ID 7A23). In both models, the structures around the FeS and SF4 ligands are of excellent quality. **b**, *Mycobacterium tuberculosis* protein FadB2 (PDB ID 6HRD). Both methods model the two Rossman folds, β1-α1-β2 and β4-α4-β5, and the α7-α11 region very well. The TopModel slightly deviates in the α2 helix. **c**, Yeast TFIIK (Kin28/Ccl1/Tfb3) complex (PDB ID 7KUE). AlphaFold2 predicts an excellent model, while TopModel places the N-lobe through the ADP binding region. This loop conformation represents another state found in the kinase structure template PDB ID 4ZSG in complex with an inhibitor.

To conclude, both TopModel and AlphaFold2 can provide high-quality enzyme structural models with, in general, very good structural features of the binding sites. In some cases, TopModel falls short in modeling loops when compared to static crystal structures. However, the exact structure of these loop regions may often be less important due to the dynamic nature of proteins.

## Conclusions

We have developed TopEnzyme, a database and framework for the structural coverage of functional enzyme space. By combining the TopEnzyme, SWISS-model, and AlphaFold DB databases, we provide the currently largest collection of enzyme structural models classified according to EC numbers. TopEnzyme provides easy access to this collection with two methods: I) A csv file containing all the metadata required for large-scale analyses. II) A treemap hosted on https://cpclab.uni-duesseldorf.de/topenzyme/ that allows the user to investigate specific enzyme functions.

With our in-house method TopModel, we added 9039 good-quality structural models, including 1297 ones of high quality. We compared a subset of these structures with AlphaFold2 models; on average, the TopScore between both models only differs by 0.04. Both methods can provide models with excellent structural features compared to experimental structures, although TopModel models sometimes differ in loop regions.

With this collection of enzyme structural models charted on functional space, researchers have access to a comprehensive and structured dataset, which should help to facilitate structure-guided investigations of specific enzymes and to develop predictive models for enzyme characteristics.

## Declarations

### Ethics approval and consent to participate

Not applicable

### Consent for publication

The publication was approved by all authors.

### Availability of data and materials

All models generated by TopModel are available in TopEnzyme at https://cpclab.uni-duesseldorf.de/topenzyme/.

## Authors’ contribution

KW developed and analyzed the database and prepared the figures. KW and HG wrote the manuscript. HG conceptualized and supervised the project, secured funding, and provided resources. All authors read and approved the final manuscript.

## Acknowledgments

We gratefully acknowledge the computing time provided by the John von Neumann Institute for Computing (NIC) on the supercomputers JUWELS and JURECA at Jülich Supercomputing Centre (JSC) (user IDs: HKF7, VSK33). We gratefully acknowledge Jonas Dittrich for the help with publishing TopEnzyme on the CPCLab website.

## References

1. The UniProt Consortium. UniProt: the universal protein knowledgebase in 2021. Nucleic Acids Res. 2021 Jan 8;49(D1):D480–9.

2. Berman H, Henrick K, Nakamura H. Announcing the worldwide Protein Data Bank. Nat Struct Biol. 2003 Dec;10(12):980.

3. Hegyi H, Gerstein M. The relationship between protein structure and function: a comprehensive survey with application to the yeast genome. J Mol Biol. 1999 Apr 23;288(1):147–64.

4. Orengo CA, Todd AE, Thornton JM. From protein structure to function. Curr Opin Struct Biol. 1999 Jun 1;9(3):374–82.

5. Nagano N, Orengo CA, Thornton JM. One Fold with Many Functions: The Evolutionary Relationships between TIM Barrel Families Based on their Sequences, Structures and Functions. J Mol Biol. 2002 Aug 30;321(5):741–65.

6. Medvedev KE, Kinch LN, Schaeffer RD, Grishin NV. Functional analysis of Rossmann-like domains reveals convergent evolution of topology and reaction pathways. PLOS Comput Biol. 2019 Dec 23;15(12):e1007569.

7. Mulnaes D, Porta N, Clemens R, Apanasenko I, Reiners J, Gremer L, et al. TopModel: Template-Based Protein Structure Prediction at Low Sequence Identity Using Top-Down Consensus and Deep Neural Networks. J Chem Theory Comput. 2020 Mar 10;16(3):1953–67.

8. Jumper J, Evans R, Pritzel A, Green T, Figurnov M, Ronneberger O, et al. Highly accurate protein structure prediction with AlphaFold. Nature. 2021 Aug;596(7873):583–9.

9. Baek M, DiMaio F, Anishchenko I, Dauparas J, Ovchinnikov S, Lee GR, et al. Accurate prediction of protein structures and interactions using a three-track neural network. Science. 2021 Aug 20;373(6557):871–6.

10. Mulnaes D, Gohlke H. TopScore: Using Deep Neural Networks and Large Diverse Data Sets for Accurate Protein Model Quality Assessment. J Chem Theory Comput. 2018 Nov 13;14(11):6117–26.

11. Varadi M, Anyango S, Deshpande M, Nair S, Natassia C, Yordanova G, et al. AlphaFold Protein Structure Database: massively expanding the structural coverage of protein-sequence space with high-accuracy models. Nucleic Acids Res. 2022 Jan 7;50(D1):D439–44.

12. Andreeva A, Howorth D, Chothia C, Kulesha E, Murzin AG. SCOP2 prototype: a new approach to protein structure mining. Nucleic Acids Res. 2014 Jan 1;42(D1):D310–4.

13. Sillitoe I, Bordin N, Dawson N, Waman VP, Ashford P, Scholes HM, et al. CATH: increased structural coverage of functional space. Nucleic Acids Res. 2021 Jan 8;49(D1):D266–73.

14. IUBMB.ORG [Internet]. IUBMB.ORG. [cited 2022 May 16]. Available from: https://iubmb.org/

15. Bienert S, Waterhouse A, de Beer Tap, Tauriello G, Studer G, Bordoli L, et al. The SWISS-MODEL Repository—new features and functionality. Nucleic Acids Res. 2017 Jan 4;45(D1):D313–9.

16. Bairoch A. The ENZYME database in 2000. Nucleic Acids Res. 2000 Jan 1;28(1):304–5.

17. Steinegger M, Söding J. MMseqs2 enables sensitive protein sequence searching for the analysis of massive data sets. Nat Biotechnol. 2017 Nov;35(11):1026–8.

18. Rost B. Twilight zone of protein sequence alignments. Protein Eng. 1999 Feb;12(2):85–94.

19. Koehl P, Levitt M. Sequence Variations within Protein Families are Linearly Related to Structural Variations. J Mol Biol. 2002 Oct 25;323(3):551–62.

20. Pearson WR. An Introduction to Sequence Similarity (“Homology”) Searching. Curr Protoc Bioinforma Ed Board Andreas Baxevanis Al. 2013 Jun;03:10.1002/0471250953.bi0301s42.

21. Craig IR, Pfleger C, Gohlke H, Essex JW, Spiegel K. Pocket-Space Maps To Identify Novel Binding-Site Conformations in Proteins. J Chem Inf Model. 2011 Oct 24;51(10):2666–79.

22. Mahajan D, Girshick R, Ramanathan V, He K, Paluri M, Li Y, et al. Exploring the Limits of Weakly Supervised Pretraining. In: Ferrari V, Hebert M, Sminchisescu C, Weiss Y, editors. Computer Vision – ECCV 2018 [Internet]. Cham: Springer International Publishing; 2018 [cited 2022 May 16]. p. 185–201. (Lecture Notes in Computer Science; vol. 11206). Available from: http://link.springer.com/10.1007/978-3-030-01216-8_12

23. Plappert M, Houthooft R, Dhariwal P, Sidor S, Chen RY, Chen X, et al. Parameter Space Noise for Exploration [Internet]. arXiv; 2018 [cited 2022 May 16]. Available from: http://arxiv.org/abs/1706.01905

24. Evans R, O’Neill M, Pritzel A, Antropova N, Senior A, Green T, et al. Protein complex prediction with AlphaFold-Multimer [Internet]. bioRxiv; 2022 [cited 2022 May 16]. p. 2021.10.04.463034. Available from: https://www.biorxiv.org/content/10.1101/2021.10.04.463034v2

25. Mirdita M, Schütze K, Moriwaki Y, Heo L, Ovchinnikov S, Steinegger M. ColabFold - Making protein folding accessible to all [Internet]. bioRxiv; 2022 [cited 2022 May 16]. p. 2021.08.15.456425. Available from: https://www.biorxiv.org/content/10.1101/2021.08.15.456425v3

26. Mariani V, Biasini M, Barbato A, Schwede T. lDDT: a local superposition-free score for comparing protein structures and models using distance difference tests. Bioinformatics. 2013 Nov 1;29(21):2722–8.

27. Soufari H, Parrot C, Kuhn L, Waltz F, Hashem Y. Specific features and assembly of the plant mitochondrial complex I revealed by cryo-EM. Nat Commun. 2020 Oct 15;11(1):5195.

28. van Eeuwen T, Li T, Kim HJ, Gorbea Colón JJ, Parker MI, Dunbrack RL, et al. Structure of TFIIK for phosphorylation of CTD of RNA polymerase II. Sci Adv. 2021 Apr 7;7(15):eabd4420.

29. Parey K, Lasham J, Mills DJ, Djurabekova A, Haapanen O, Yoga EG, et al. High-resolution structure and dynamics of mitochondrial complex I-Insights into the proton pumping mechanism. Sci Adv. 2021 Nov 12;7(46):eabj3221.

30. Cox J a. G, Taylor RC, Brown AK, Attoe S, Besra GS, Fütterer K. Crystal structure of Mycobacterium tuberculosis FadB2 implicated in mycobacterial β-oxidation. Acta Crystallogr Sect Struct Biol. 2019 Jan 1;75(1):101–8.

31. Luque I, Freire E. Structural stability of binding sites: consequences for binding affinity and allosteric effects. Proteins. 2000;Suppl 4:63–71.

