## Supporting Information for "TopEnzyme: A framework and database for structural coverage of the functional enzyme space"

Author ORCID:

*Corresponding Author:

Holger Gohlke

Phone: (+49) 2461 61 85550

### Table of Contents

#### Supplemental Data

**Data S1.** Treemap visualisation of TopEnzyme. See file TopEnzyme.html or https://cpclab.uni-duesseldorf.de/topenzyme/ ………………………………………………4

**Data S2.** Csv file containing the meta-data for each UniprotAC identifier. See file TopEnzyme.csv or obtain it from https://cpclab.uni-duesseldorf.de/topenzyme/ ………..4

#### Supplemental Figures

**Figure S1.** TopModel models generated without and with refinement procedure. The refined models were created using the TopModel webserver (https://cpclab.uni-duesseldorf.de/topsuite/topmodel.php). Ten enzyme structures were randomly selected from each enzyme mainclass for the full modeling procedure. The average unsigned difference between the TopScore values is 0.06, with models of better quality obtained after refinement. ……………………………………………………………………………...5

**Figure S2.** pLDDT from AlphaFold2 against (1 – TopScore) (1 - TopScore was linearly rescaled to lDDT range [0-100]) for all 2419 AlphaFold2 structural models. The red line is the linear correlation between both scores ($p < 0.001$, $R^2 = 0.59$). The average unsigned difference is 16 lDDT. With respect to data points in the bottom right corner, see the performance of pLDDT and (1 – TopScore) against lDDT depicted in Figure S3. ………...6

**Figure S3.** pLDDT and scaled (1 – TopScore) computed for AlphaFold2-predicted structural models against the lDDT determined by comparison to 35 experimental structures. The experimental structures were recently deposited in the PDB and were not seen during training by AlphaFold2 or TopModel. The pLDDT rates the model better than lDDT, whereas the (1 – TopScore) undervalues the model compared to lDDT. …………..7

**Supplemental Data**

**Data S1. TopEnzyme.html**

Treemap visualisation of TopEnzyme. See file TopEnzyme.html or https://cpclab.uni-duesseldorf.de/topenzyme/

**Data S2. TopEnzyme.csv**

Csv file containing the meta-data for each UniprotAC identifier. See file TopEnzyme.csv or obtain it from https://cpclab.uni-duesseldorf.de/topenzyme/

**Supplemental Figures**

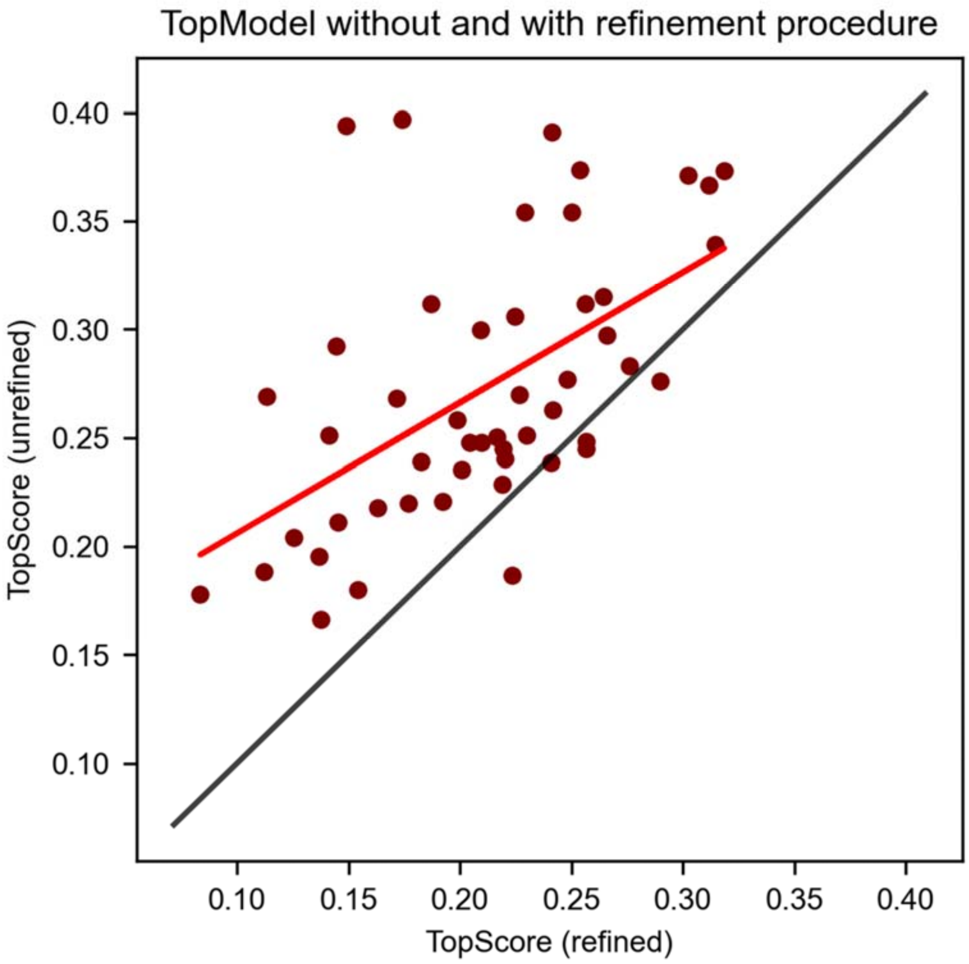


**Figure S1. TopModel without and with refinement procedure** TopModel models generated without and with refinement procedure. The refined models were created using the TopModel webserver (https://cpclab.uni-duesseldorf.de/topsuite/topmodel.php). Ten enzyme structures were randomly selected from each enzyme mainclass for the full modeling procedure. The average unsigned difference between the TopScore values is 0.06, with models of better quality obtained after refinement.

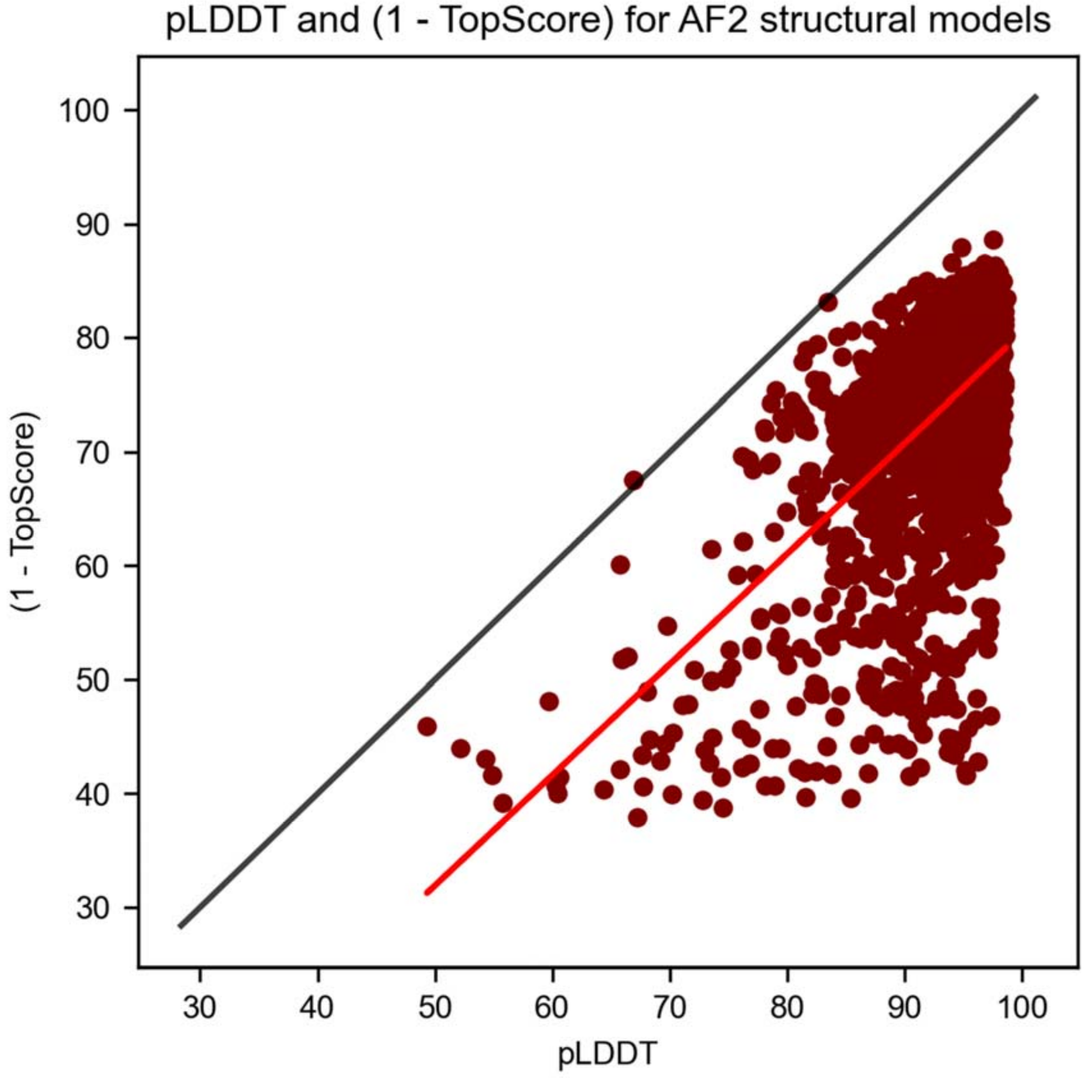


**Figure S2. pLDDT and (1 – TopScore) for AlphaFold2 structural models.** pLDDT from AlphaFold2 against (1 – TopScore) (1 - TopScore was linearly rescaled to lDDT range [0-100]) for all 2419 AlphaFold2 structural models. The red line is the linear correlation between both scores ($p < 0.001$, $R^2 = 0.59$). The average unsigned difference is 16 lDDT. With respect to data points in the bottom right corner, see the performance of pLDDT and (1 – TopScore) against lDDT depicted in Figure S3.

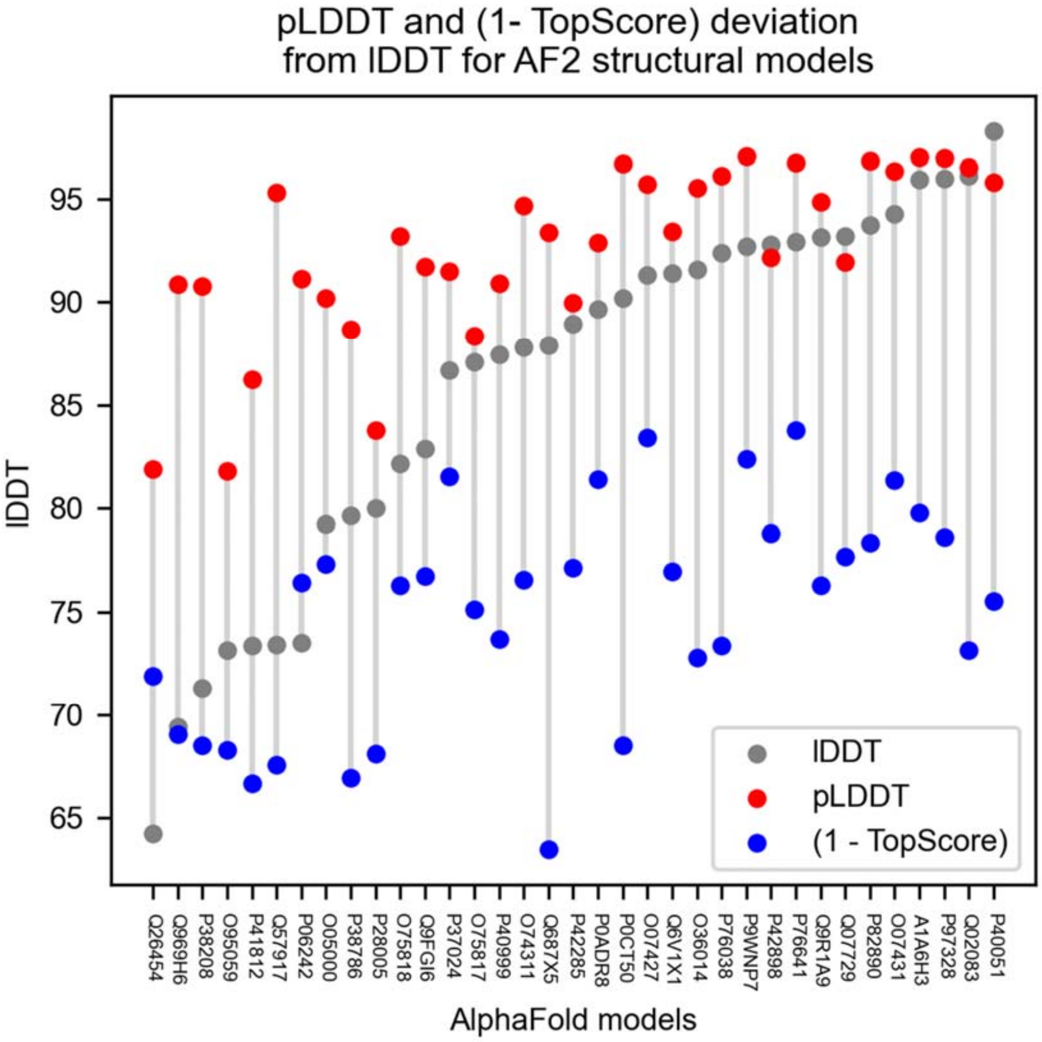


**Figure S3. pLDDT and (1 – TopScore) deviations from lDDT for AlphaFold2 structural models.**

pLDDT and scaled (1 – TopScore) computed for AlphaFold2-predicted structural models against the lDDT determined by comparison to 35 experimental structures. The experimental structures were recently deposited in the PDB and were not seen during training by AlphaFold2 or TopModel. The pLDDT rates the model better than lDDT, whereas the (1 – TopScore) undervalues the model compared to lDDT.
